# Familial prion disease-related mutation E196K displays a novel amyloid fibril structure revealed by cryo-EM

**DOI:** 10.1101/2021.02.18.431846

**Authors:** Li-Qiang Wang, Kun Zhao, Han-Ye Yuan, Xiang-Ning Li, Hai-Bin Dang, Yeyang Ma, Qiang Wang, Chen Wang, Yunpeng Sun, Jie Chen, Dan Li, Delin Zhang, Ping Yin, Cong Liu, Yi Liang

## Abstract

Prion diseases are caused by the conformational conversion of prion protein (PrP) from its cellular form (PrP^C^) into a protease-resistant, aggregated form (PrP^Sc^). 42 different familial mutations were identified in human PrP, which lead to genetic prion diseases with distinct clinical syndromes. Here we report cryo-EM structure of an amyloid fibril formed by full-length human PrP with E196K mutation, a familial Creutzfeldt-Jakob disease-related mutation. This mutation disrupts key interactions in wild-type PrP fibril and results in a rearrangement of the overall structure, forming an amyloid fibril with a conformation distinct from wild-type PrP fibril. The E196K fibril consists of two protofibrils intertwined into a left-handed helix. Each subunit forms five β-strands stabilized by a disulfide bond and an unusual hydrophilic cavity. Two pairs of amino acids (Lys194 and Glu207; Lys196 and Glu200) from opposing subunits form four salt bridges to stabilize the zigzag interface of the two protofibrils. Furthermore, the E196K fibril exhibits a significantly lower conformational stability and protease resistance activity than the wild-type fibril. Our results provide direct structural evidences of the diverse mammalian prion strains and fibril polymorphism of PrP, and highlight the importance of familial mutations in determining the different prion strains.

## INTRODUCTION

Prions, which mean ‘protein infectious agents’ and convert between structurally and functionally distinct states, were originally isolated and described by S. B. Prusiner (*1−3*). Prion diseases are infectious, fatal neurodegenerative diseases primarily caused by the conformational conversion of prion protein (PrP) from its cellular form (PrP^C^) into a protease-resistant, aggregated form (PrP^Sc^) in humans, cattle, sheep and cervid species (*1−12*). 42 different mutations in the prion protein gene (*PRNP*) were identified to cause a variety of genetic prion diseases including familial Creutzfeldt−Jakob disease (CJD), Gerstmann−Sträussler−Scheinker disease (GSS) and fatal familial insomnia (FFI) (*1, 4, 6, 10*). These disease-related mutations can form different strains with distinct conformations, and display distinct clinical syndromes with different incubation times (*13−20*). Most of the mutations do not cause large conformational changes in PrP^C^, whose structure features a folded C-terminal globular domain containing three α-helices and two very short antiparallel β-sheets accompanied by a largely disordered N-terminal tail (*5, 10, 21*). Instead, the mutations were reported to induce spontaneous generation of PrP^Sc^ in the brain of patients with genetic prion diseases^10^ and in the brain of transgenic mouse models with the mutations (*22, 23*). Since prions were discovered in 1982 (*1*), great efforts have been dedicated to unravel the mysteries of the atomic structure of prion (*7*, *8*, *11*, *20*, *24−33*) and prion strains (*13−20*). Recently, we reported a cryo-EM structure of the amyloid fibril formed by full-length wild-type human PrP featuring a parallel in-register intermolecular β-sheet architecture (*33*), which provides structural insights into the conversion from α-helix-dominant PrP^C^ to β-sheet-rich PrP^Sc^.

The first reported patient with E196K mutation in the *PRNP* gene, a familial CJD-related mutation, was a French 69-year-old woman, who died 1 year after the first recorded symptoms (*34*). The patients carrying this mutation have an average age at onset of 71.2 ± 5.9 years old (*n* = 15) and feature rapid disease progression and short disease duration (6.6 ± 3.5 months) (*35, 36*). Previous studies have shown that E196K mutation decreases the thermal stability of PrP^C^ and thus increases the propensity for PrP amyloid formation (*37*). The atomic structure of wild-type PrP fibrils has shown that several familial mutations including K194E, E196K and E211Q may break salt bridges essential for forming the protofilament interface in wild-type PrP fibrils and thus disrupt PrP fibril structure (*33*). Whether E196K induces a new fibril fold is unknown.

Here we prepared homogeneous amyloid fibrils in vitro from recombinant, full-length human E196K PrP^C^, and determined the atomic structure by using cryo-EM. While the infectivity of these fibrils remains to be established, the structural features provide insights into the mechanism how familial mutations drive formation of different prion strains.

## RESULTS

### E196K forms amyloid fibrils morphologically distinct from wild-type PrP

We produced amyloid fibrils from recombinant, full-length human E196K PrP^C^ (residues 23−231) overexpressed in *Escherichia coli*, by incubating the purified protein in 20 mM Tris-HCl buffer (pH 7.4) containing 2 M guanidine hydrochloride and shaking at 37 °C for 7−9 h. E196K fibrils were dialyzed against NaAc buffer, purified by ultracentrifugation, resuspended in NaAc buffer and examined by transmission electron microscopy (TEM) without further treatment.

Negative-staining TEM imaging showed that E196K PrP^C^ formed homogeneous and unbranched fibrils (Fig. 1A), similar to wild-type PrP fibrils formed at the same conditions (Fig. 1B). The E196K fibril is composed of two protofibrils intertwined into a left-handed helix, with a fibril width of 22 nm and a helical pitch of 272 nm (Fig. 1A). By comparison, the wild-type fibril is also composed of two protofibrils intertwined into a left-handed helix, but with a fibril width of 26 nm and a short helical pitch of 153.6 nm (Fig. 1B). Thus, E196K formed an amyloid fibril with a morphology distinct from wild-type PrP.

**Fig. 1.**
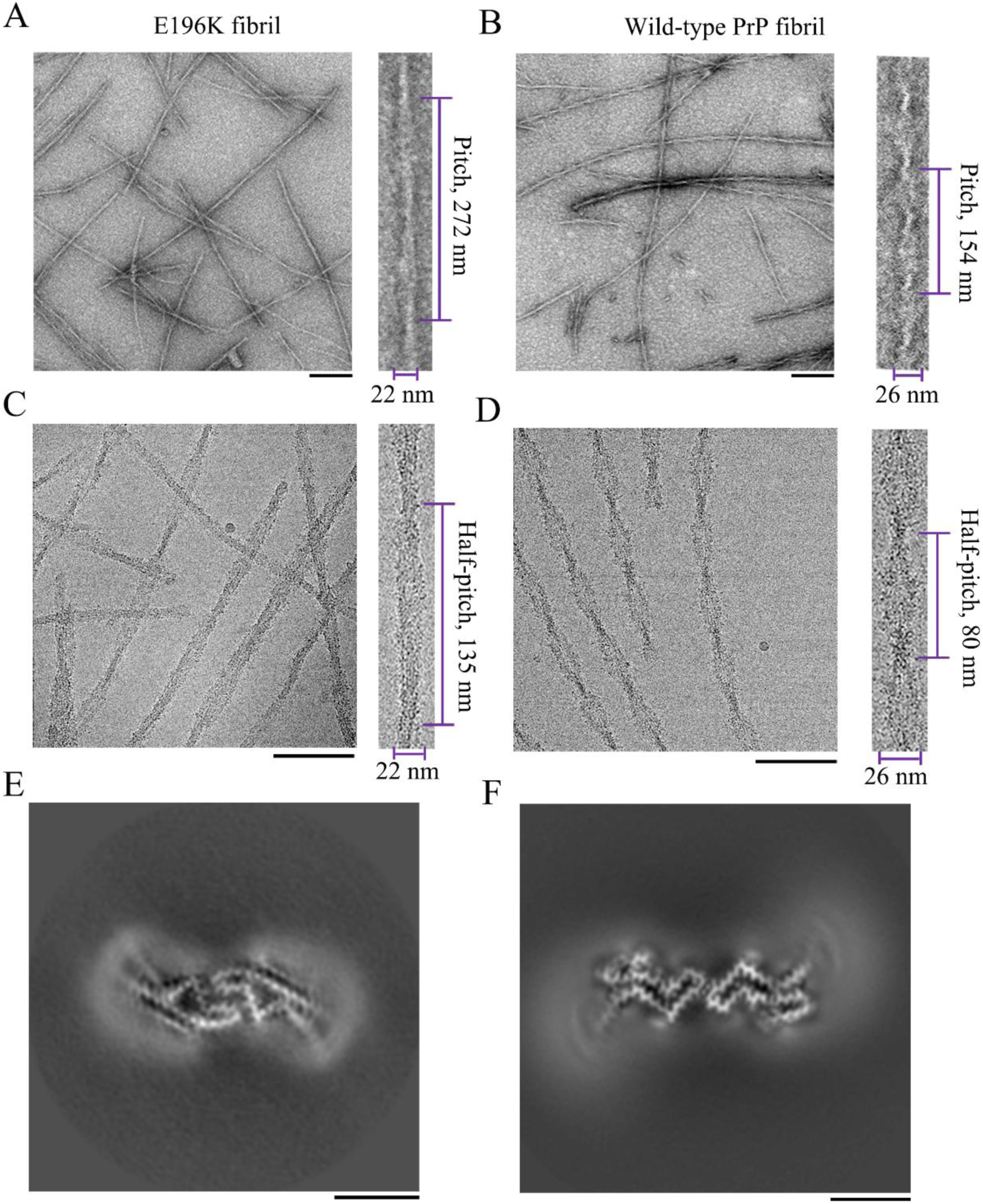
Comparison of the images of E196K fibril and the wild-type fibril. (**A and B**) Negative-staining TEM images of amyloid fibrils from full-length wild-type human PrP (**B**) and its E196K variant (**A**). Enlarged section of **A** (right) or **B** (right) showing two protofibrils intertwined into a left-handed helix, with a fibril width of 22 nm (**A**) or 26 nm (**B**) and a helical pitch of 272 nm (**A**) or 153.6 nm (**B**). The scale bars represent 200 nm. (**C** and **D**) Raw cryo-EM images of amyloid fibrils from E196K (**C**) and its wild-type form (**D**). Enlarged section of **C** (right) or **D** (right) showing two protofibrils intertwined into a left-handed helix, with a fibril width of 22 nm (**C**) or 26 nm (**D**) and a half-helical pitch of 135 nm (**C**) or 80 nm (**D**). The scale bars represent 100 nm. E196K formed amyloid fibril with a morphology distinct from the one formed by wild-type PrP. (**E** and **F**) Cross-sectional view of the 3D map of E196K fibril (**E**) showing two protofibrils also forming a dimer but with a conformation distinct from the wild-type fibril (**F**). Scale bars, 5 nm.

Congo red binding assays showed a red shift of the maximum absorbance, from 490 to 550 nm, in the presence of E196K fibrils (fig. S1A), which is typical of amyloid fibrils (*33*). This is similar to wild-type PrP fibrils formed at the same conditions (*33*). Proteinase K digestion of the E196K fibrils generated a predominant band with an apparent molecular weight of 14−15 kDa (fig. S1C). This is different from wild-type PrP fibrils formed at the same conditions, which generated a predominant band with an apparent molecular weight of 15−16 kDa (*33*). Furthermore, at the same protease:PrP molar ratios, the optical density of the 14−15-kDa band in the E196K fibrils was remarkably weaker than that of the 15−16-kDa band in the wild-type fibrils (fig. S1, B and C). In addition, we observed two other bands with apparent molecular weights of 12 and 10 kDa after proteinase K digestion of the E196K fibrils (fig. S1C), which is similar to wild-type PrP fibrils (fig. S1B). Together, the data showed that E196K fibril exhibits a distinct morphology with weaker protease resistance activity compared to the wild-type fibril.

### Cryo-EM structure of E196K fibrils

We determined the atomic structure of the E196K amyloid fibrils by cryo-EM (Table 1). The cryo-EM micrographs and two-dimensional (2D) class average images show that the E196K fibril is composed of two protofibrils intertwined into a left-handed helix (Fig. 1C and fig. S2A), with a helical pitch longer than the wild-type fibril (Fig. 1, C and D). Furthermore, two protofilaments in each E196K fibril is arranged in a staggered manner (fig. S2, B and C), similar to wild-type PrP fibrils formed at the same conditions (*33*). The fibrils are morphologically homogeneous, showing a fibril width of 22 nm (Fig. 1C and fig. S2A). This is narrower than wild-type PrP fibrils (*33*) but similar in size to previously described ex vivo, infectious PrP^Sc^ fibrils, which showed width of ∼20 nm based on negative staining on TEM (*31, 38*).

**Table 1.**
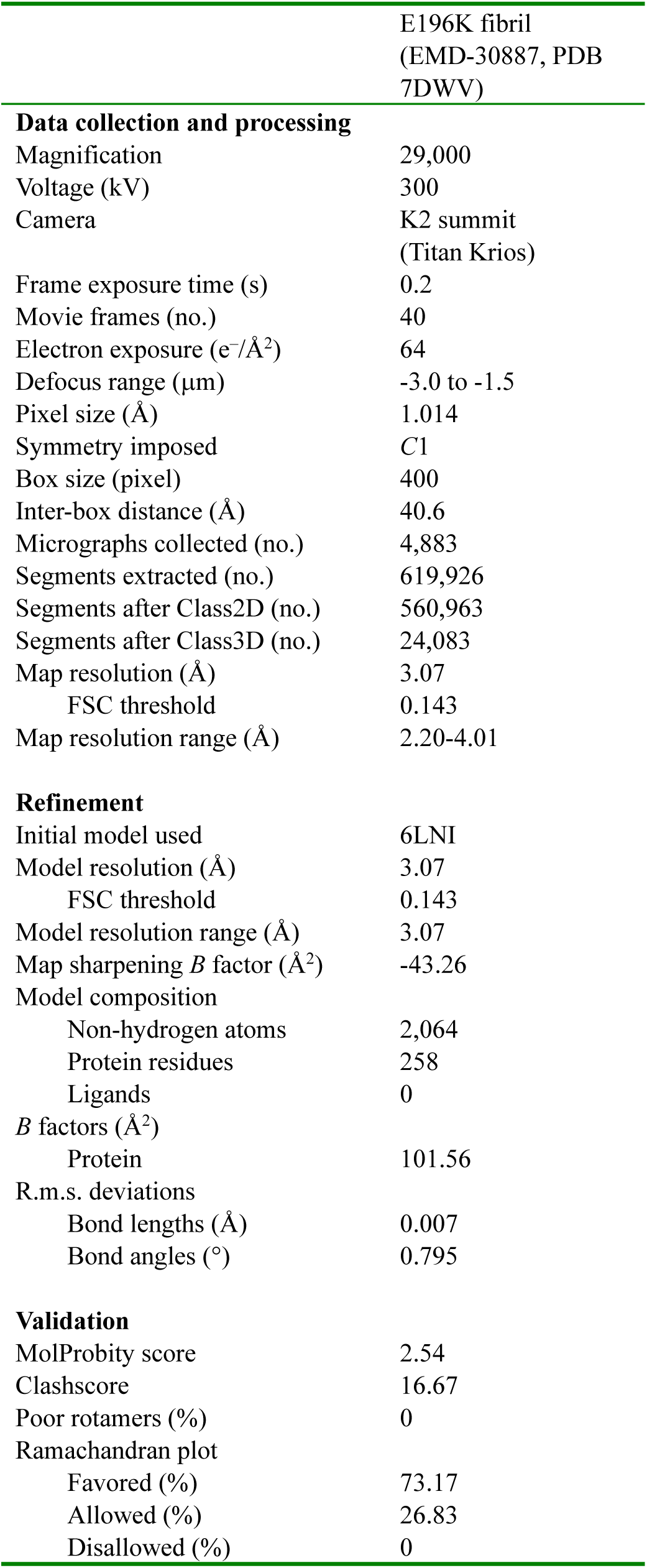
Cryo-EM data collection, refinement and validation statistics

Using helical reconstruction in RELION3.1 (*39*), we determined a density map of the ordered core of E196K fibrils, with an overall resolution of 3.07 Å, which features well-resolved side-chain densities and clearly separated β-strands along the helical axis (Fig. 1E and fig. S3). The three-dimensional (3D) map showed two protofibrils in the E196K fibril intertwined into a left-handed helix, with a half-helical pitch of 126.4 nm (Fig. 2A) which is remarkably longer than that of the wild-type fibril (*33*). The width of the fibril core is ∼11 nm (Fig. 2A). This is narrower than wild-type PrP fibrils formed at the same conditions (*33*) but similar in size to previously described ex vivo, infectious PrP^Sc^ fibrils, which showed width of ∼10 nm based on cryo-EM images (*28*). Cross-sectional view of the 3D map of E196K fibril (Fig. 1E) shows two protofibrils conformationally distinct from those of wild-type PrP fibril (Fig. 1F). The protofilaments in the E196K fibril form a dimer with a screw symmetry of approximately 2_1_ (Fig. 2, B−D). The subunits in each E196K protofibril stack along the fibril axis with a helical rise of 4.82 Å and twist of −0.68° (Fig. 2E), and the subunits in two protofibrils stack along the fibril axis with a helical rise of 2.41 Å (Fig. 2F).

**Fig. 2.**
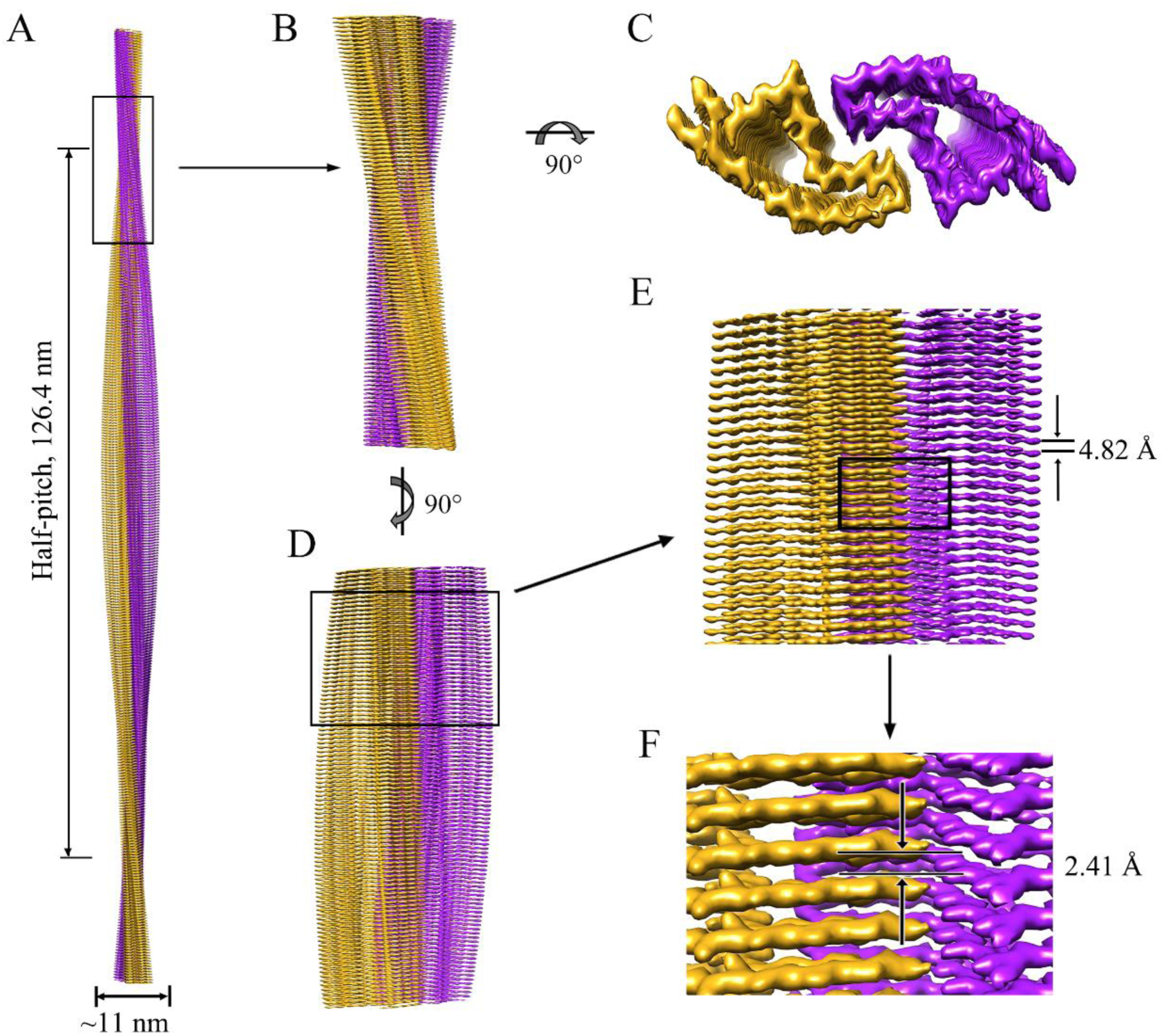
Cryo-EM structure of E196K fibrils. (**A**) 3D map showing two protofibrils intertwined into a left-handed helix, with a fibril core width of ∼11 nm and a half-helical pitch of 126.4 nm. The two intertwined protofibrils are colored purple and golden, respectively. (**B**) Enlarged section showing a side view of the density map. (**C**) Top view of the density map. (**D**) Another side view of the density map after 90° rotation of **B** along the fibril axis. (**E**) Close-up view of the density map in **D** showing that the subunits in each protofibril stack along the fibril axis with a helical rise of 4.82 Å. (**F**) Close-up view of the density map in **E** showing that the subunits in two protofibrils stack along the fibril axis with a helical rise of 2.41 Å.

We unambiguously built a E196K fibril model comprising residues 175−217 at 3.07 Å (Fig. 3) and an unmasked E196K fibril model comprising residues 171−222 at 3.59 Å (fig. S4), both of which are slightly shorter than the wild-type PrP fibril core comprising residues 170−229 (*33*). Side chain densities for most residues in E196K fibril core had high local resolution (2.20−3.28 Å) (fig. S4, A to C, and Fig. 3, A and B). The exterior of the E196K fibril core is mostly hydrophilic. The fibril core is stabilized by an intramolecular disulfide bond between Cys179 and Cys214 (Fig. 3, A and B, and figs. S4, B and C, and S5, A and B) (also present in both PrP^C^ and PrP^Sc^) (*24, 27, 28, 30*) and a hydrogen bond between His177 and Thr216 (fig. S5, A and B) in each monomer. In sharp comparison to the highly hydrophobic cavity observed in the wild-type fibril, E196K fibril core is composed of an unusual hydrophilic cavity in each subunit (Fig. 3, B and G, and fig. S4C) which may decrease the fibril stability of E196K.

We observed two unidentified densities flanking the two protofibrils in E196K fibril, termed two islands (fig. S4 and Fig. 3A), which are reminiscent of those islands observed in the structures of filaments formed by α-synuclein hereditary disease mutant H50Q (*40*). Each island, probably comprising residues 136−142 from E196K and forming a β-strand (β0), is located on the opposing side of hydrophobic side chains of Val180, Ile182 and Ile184 in each monomer (fig. S4, B to D). The side chains of Val180, Ile182, Ile184, Pro137, Ile139 and Phe141 forms a hydrophobic steric zipper-like interface, thereby stabilizing the E196K fibrils. The presence of two islands represents one of the major structural difference between E196K and wild-type fibrils.

**Fig. 3.**
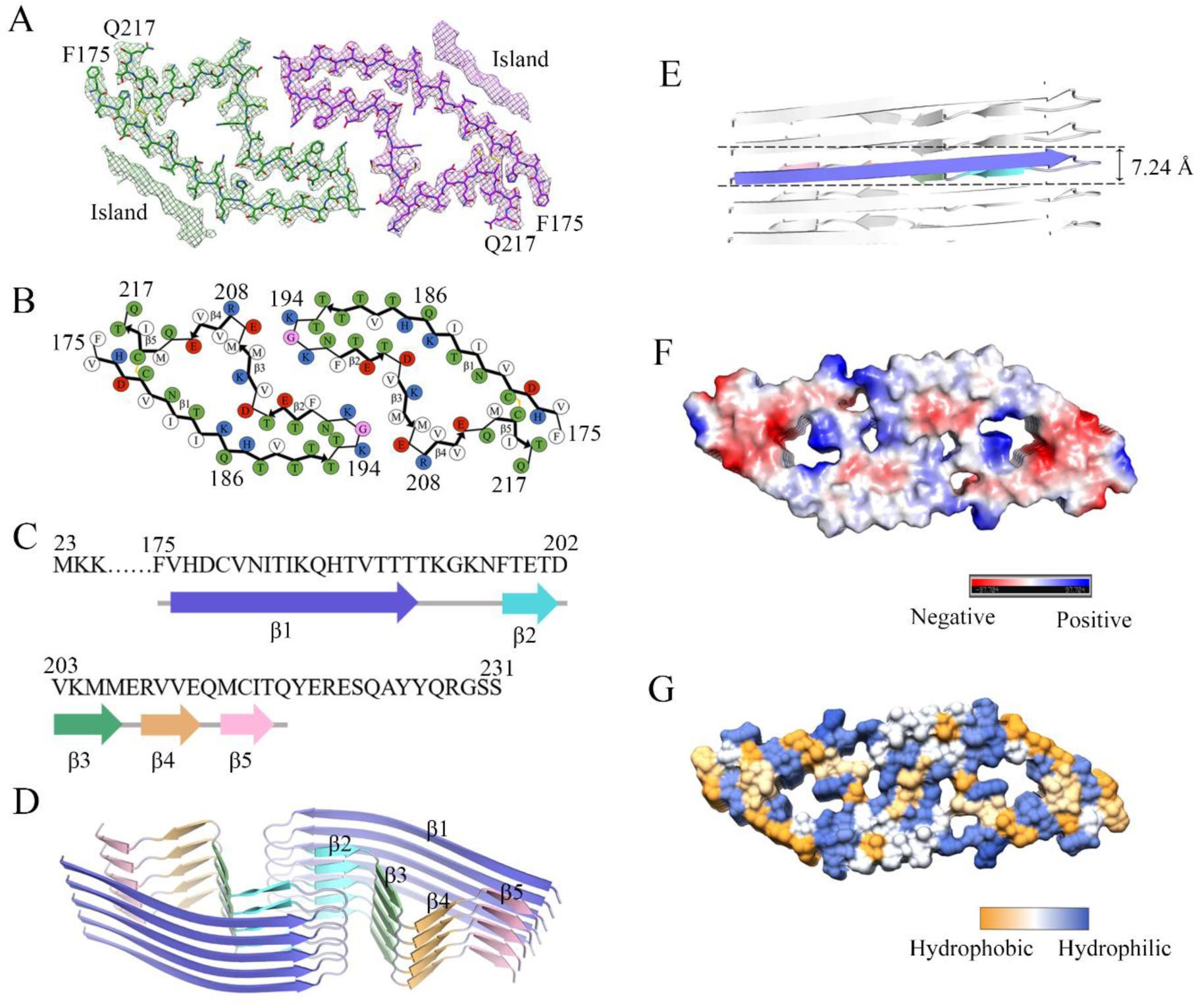
Atomic structure of E196K fibrils. (**A**) Cryo-EM map of E196K fibrils with the atomic model overlaid. Two identical densities (termed two islands) flanking the two protofibrils, which are colored green and purple, respectively. Each island is located on the opposing side of hydrophobic side chains of Val180, Ile182 and Ile184 in each monomer. (**B**) Schematic view of E196K fibril core. Residues are colored as follows: white, hydrophobic; green, polar; red and blue, negatively and positively charged, respectively; magenta, glycine. β-strands are indicated with bold lines. E196K fibrils are stabilized by a disulfide bond (yellow line) formed between Cys179 and Cys214 in each monomer, also visible in panel **A**. (**C**) Sequence of the fibril core comprising residues 175−217 from E196K, with the observed five β-strands colored blue (β1), cyan (β2), green (β3), orange (β4) and pink (β5). The dashed line corresponds to residues (23−174) not modeled in the cryo-EM density. (**D**) Ribbon representation of the structure of an E196K fibril core containing five molecular layers and a dimer. (**E**) As in **D**, but viewed perpendicular to the helical axis, revealing that the height of one layer of the subunit along the helical axis is 7.24 Å. (**F**) Electrostatic surface representation of the structure of an E196K fibril core containing five molecular layers and a dimer. (**G**) Hydrophobic surface representation of the structure of an E196K fibril core as in **D**. (**F** and **G**) Two pairs of amino acids (Lys194 and Glu207; Lys196 and Glu200) from opposing subunits form four salt bridges at the zigzag interface of the two protofibrils. The surface of two opposing subunits is shown according to the electrostatic properties (F) or the hydrophobicity (**G**) of the residues.

Five β-strands (β1−β5) and six β-strands (β0−β5) are present in the E196K fibril core structure at 3.07 Å (Fig. 3, B to D) and an unmasked structure at 3.59 Å (fig. S4C), respectively. The E196K fibril core features a compact fold containing a long β-strand (β1) and four short β-strands (β2−β5). This is different from a previously observed wild-type PrP fibril core, which contains six short β-strands (β1−β6) (*33*). The height of one layer of the subunit along the helical axis is 7.24 Å (Fig. 3E). A U-turn between β1 and β2 containing residues ^193^TKGKN^197^ enables antiparallel cross-β packing of the first part of β1 against β2, and a disulfide bond between Cys179 in β1 and Cys214 in β5 enables antiparallel cross-β packing of the last part of β1 against β5 (Fig. 3, B and D, and fig. S4C).

In the E196K fibril, two pairs of amino acids (Lys194 and Glu207; Lys196 and Glu200) from opposing subunits form four salt bridges at the zigzag interface of the two protofibrils (Figs. 2C and 3, A, F and G, and fig. S4, B and C). The interfaces in the E196K fibril feature mixed compositions of hydrophilic and hydrophobic side chains (Fig. 3, B and G, and fig. S4C). They are reminiscent of those interfaces observed in the structures of Tau filaments extracted from the brains of patients with corticobasal degeneration (*41*) and amyloid fibrils formed by the S20G mutation in human amylin (*42*). In the wild-type PrP fibril, however, Lys194 and Glu196 from opposing subunits form two salt bridges that create a hydrophilic cavity at the interface of the two protofibrils with two additional unidentified densities (*33*).

We then compare protofilament interfaces of E196K and wild-type fibrils in detail (Fig. 4). E196K fibril has a long interface comprising residues 194−208, but the wild-type fibril features a very short interface comprising only three residues 194−196 (Fig. 4A). Four pairs of intermolecular salt bridge formed by Glu207 & Lys194 and Glu200 & Lys196 (Fig. 4, B to E) are identified in the zigzag interface between E196K protofibrils. In contrast, two intermolecular salt bridge between Lys194 and Glu196 are formed at the dimer interface between wild-type PrP protofibrils (Fig. 4, F and G). Thus, the E196K mutation disrupts the key salt bridges in wild-type PrP fibrils and results in a rearrangement of the overall structure, forming an amyloid fibril with a conformation distinct from the wild-type fibril (Figs. 4 and 5 and fig. S5).

**Fig. 4.**
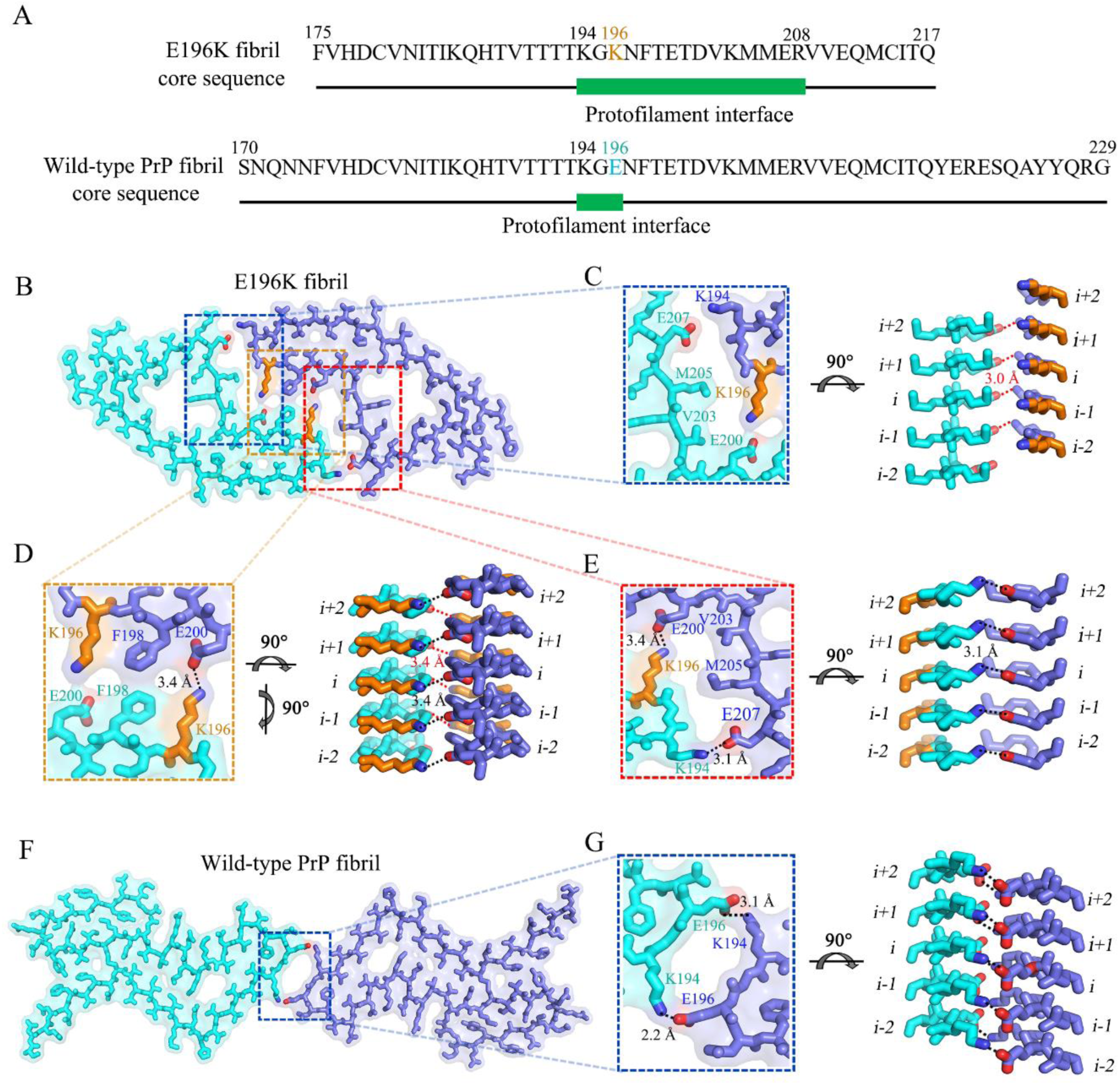
Comparison of protofilament interfaces of E196K and wild-type fibrils. (**A**) The primary sequences of E196K fibril core and the wild-type fibril core. The green bar marks the region of the protofilament interface. Lys196 in E196K variant and Glu196 in wild-type PrP are highlighted in orange and cyan, respectively. E196K fibril has a long interface comprising residues 194−208, but the wild-type fibril has a very short interface comprising only three residues 194−196. (**B**) A space-filled model overlaid onto a stick representation of E196K fibril in which one protofibril is shown in cyan and another in blue. Lys/Glu pairs that form salt bridges are highlighted in red (oxygen atoms in Glu), blue (nitrogen atom in Lys) and orange (Lys196), and the dimer interface is magnified in **C**, **D** and **E**. (**C**) A magnified top view of the top region of the zigzag interface between E196K protofibrils, where two pairs of amino acids (Glu207 and Lys194; Glu200 and Lys196) from opposing subunits form two salt bridges. A side view (right) highlighting a salt bridge between Glu207 (*i*) and Lys194 from its opposing adjacent subunit (*i-1*), with a distance of 3.0 Å (red). (**D**) A magnified top view of the middle region of the interface between E196K protofibrils, where two pairs of amino acids (Glu200 and Lys196; Lys196 and Glu200) from opposing subunits form two salt bridges. A side view (right) highlighting a salt bridge between Glu200 (*i*) and Lys196 from its opposing adjacent subunit (*i-1*), with a distance of 3.4 Å (red), and another salt bridge between Lys196 (*i*) and Glu200 from its opposing subunit (*i*), also with a distance of 3.4 Å (black). (**E**) A magnified top view of the bottom region of the interface between E196K protofibrils, where two pairs of amino acids (Lys196 and Glu200; Lys194 and Glu207) from opposing subunits form two salt bridges. A side view (right) highlighting a salt bridge between Lys194 (*i*) and Glu207 from its opposing subunit (*i*), with a distance of 3.1 Å (black). (**F)** A space-filled model overlaid onto a stick representation of wild-type PrP fibril (PDB 6LNI) (*33*) in which one protofibril is shown in cyan and another in blue, and the protofilament interface is magnified in **G**. (F) A magnified top view of the dimer interface between wild-type PrP protofibrils, where two pairs of amino acids (Lys194 and Glu196; Glu196 and Lys194) from opposing subunits form two salt bridges. A side view (right) highlighting a salt bridge between Lys194 (*i*) and Glu196 from its opposing subunit (*i*), with a distance of 2.2 Å (black), and another salt bridge between Glu196 (*i*) and Lys194 from its opposing subunit (*i*), with a distance of 3.1 Å (black).

**Fig. 5.**
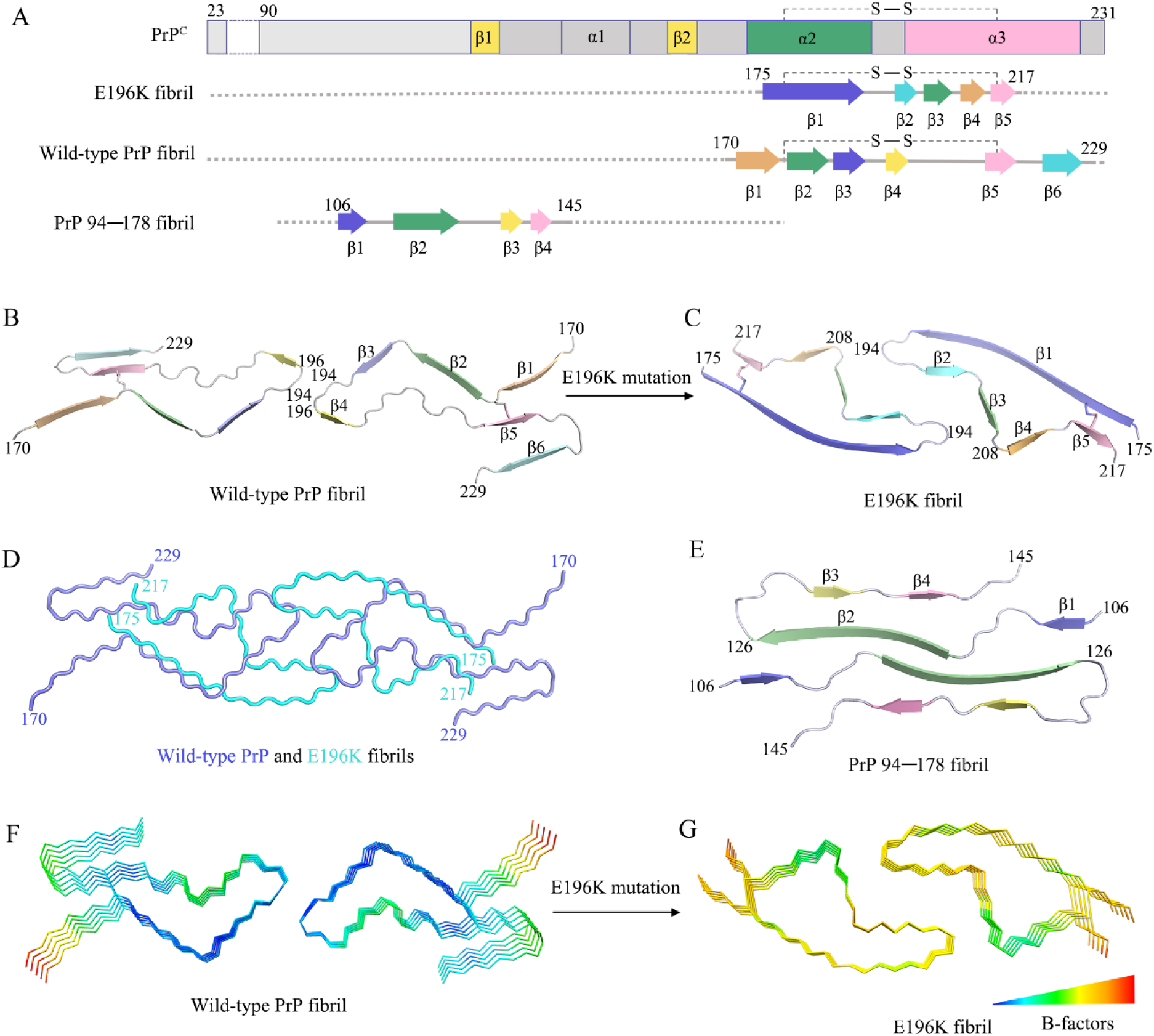
Comparison of the structures of PrP^C^, E196K fibril, wild-type PrP fibril and PrP_94−178_ fibril. (**A**) Sequence alignment of the full-length wild-type human PrP^C^ (23−231) monomer with two short β-sheets, three α-helices and a single disulfide bond between Cys179 in α2 and Cys214 in α3 (PDB 1QLX) (*21*), the E196K fibril core comprising residues 175−217 from E196K with the observed five β-sheets and one disulfide bond between Cys179 in β1 and Cys214 in β5, the wild-type PrP fibril core comprising residues 170−229 from wild-type PrP with the observed six β-sheets and one disulfide bond between Cys179 in a loop (linking β1 to β2) and Cys214 in β5 (PDB 6LNI) (*33*) and a human PrP fragment 94−178 fibril core comprising residues 106−145 with the observed four β-sheets (PDB 6UUR) (*32*). S – S denotes a single disulfide bond. Dashed box (**A**) corresponds to residues (23−174) for the E196K fibril, residues (23−169) for the wild-type fibril and residues (94−105)/ (146−178) for the PrP 94−178 fibril, respectively, which were not modeled in the cryo-EM density. (**B** and **C**) Ribbon representation of the structures of a wild-type PrP fibril core (PDB 6LNI) (*33*) (**B**) and an E196K PrP fibril core (**C**), both of which contain one molecular layer and a dimer with a single disulfide bond in each monomer. (**D**) Overlay of the structures of a wild-type PrP fibril core (blue) (PDB 6LNI) (*33*) and an E196K PrP fibril core (cyan). The E196K mutation disrupts important interactions in PrP fibrils and results in a rearrangement of the overall structure, forming an amyloid fibril with a dimer interface and a conformation distinct from the wild-type fibril. (**E**) Ribbon representation of the structure of a PrP_94-178_ fibril core containing one molecular layer and a dimer (PDB 6UUR) (*32*). (**F** and **G**) B-factor representation of the structures of a wild-type PrP fibril core (PDB 6LNI) (*33*) (**F**) and an E196K PrP fibril core (**G**), both of which contain five molecular layers and a dimer. The C^α^ atoms of the fibrils are colored according to B-factors, ranging from blue (69 Å^2^, lowest) to red (121 Å^2^, highest). The local region comprising residues 180−194 has remarkably higher flexibility in E196K fibril (yellow) (**G**) compared to that in the wild-type fibril (blue) (**F**).

### The E196K mutation significantly decreases the conformational stability of PrP fibrils

Compared to the wild-type fibril, E196K fibril features a smaller and distinct fibril core with an unusual hydrophilic cavity in the center. We next examined whether E196K fibril exhibits distinct conformational stability from the wild-type fibril. Chemical and/or thermal denaturation was widely used to evaluate the conformational stability of proteinase resistant PrP fibrils (*19, 43, 44*). A strong chaotropic salt, guanidine thiocyanate, was used in our denaturation assay (fig. S6A). The *C*_1⁄2_ value of wild-type PrP fibril is 2.00 ± 0.03 M (fig. S6B) which is consistent with previous data (*42*). Notably, the *C*_1⁄2_ value of E196K fibrils is 1.54 ± 0.06 M (fig. S6B) which is significantly lower than the wild-type fibril, suggesting that E196 fibril is less stable that the wild type.

To validate the stability differences between E196K and wild-type fibrils, we further measured their thermostabilities. The PrP fibrils were incubated with 6% SDS under a thermal gradient from 25−100 °C, and the soluble PrP disassembled from fibrils was measured by SDS-PAGE. The results showed that the melting temperature (*T*_m_) value of E196K fibrils is ∼75 °C, substantially lower than that of the wild-type fibrils (∼95 °C) (fig. S6, C and D). Together, these results demonstrate that the E196K fibril has a significantly lower conformational stability compared to the wild-type fibril.

## DISCUSSION

We compared the secondary structures of PrP^C^, E196K fibril, wild-type fibril and PrP_94−178_ fibril (Fig. 5). Strikingly, the PrP molecule adopts largely distinctive secondary structures in four different PrP structures, highlighting the high structural polymorphs of PrP in soluble and fibrillar forms. The human PrP^C^ contains three α-helices, two very short antiparallel β-sheets and a single disulfide bond between Cys179 in α2 and Cys214 in α3 (*5, 10, 21*) (Fig. 5A). Once it folds into its fibrillar form, the PrP subunit undergoes a totally conformational rearrangement. The wild-type PrP fibril core contains six short β-sheets (β1−β6), a very short interface comprising only three residues 194−196 and one disulfide bond between Cys179 in a loop (linking β1 to β2) and Cys214 in β5 (*33*) (Fig. 5, A and B). In contrast, the E196K fibril core contains a long β-sheet (β1), four short β-sheets (β2−β5), a long interface comprising residues 194−208 and one disulfide bond between Cys179 in β1 and Cys214 in β5 (Fig. 5, A and C), with a higher β-sheet content (77%) than in the wild-type fibril core (55%) (Fig. 5, A to C). These results provide structural evidence that different prion strains have distinct conformations, which may underscore a pivotal role of the specific structural features in driving the disease phenotype.

Conformational stabilities of PrP^Sc^ and PrP fibrils assessed by measuring their resistance to chemical and/or thermal denaturation are used to probe the structural differences between prion strains (*13, 16, 19, 20, 43, 44*). However, the structural basis of the stability differences between different strains remains to be established (*19, 20, 43, 44*). Our structure data revealed the key structural differences between E196K and wild-type fibrils including (1) hydrophilic cavity VS. hydrophobic cavity; (2) a mixed hydrophilic and hydrophobic protofibril interface VS. a short hydrophilic protofibril interface. This may help to understand how a single point mutation may lead to formation of a new prion strain with largely distinct properties.

Asn181 and Asn197 were previously reported to be glycosylated in wild-type PrP^Sc^ (*24, 27, 30*). The recombinant PrPs we used are non-glycosylated. But, in our wild-type PrP fibril model, the side chains of the two Asn residues appear well exposed to solvent (*33*) and thus are able to sterically accommodate bulky, N-linked glycans as shown by recent molecular dynamics simulations (*45, 46*). Intriguingly, in the E196K fibril structure, the side chains of Asn181 and Asn197 are buried in the interior of a hydrophilic core (Fig. 3, A and B). Notably, the side chain of Asn181 in the E196K fibril fold appears well exposed to an open area of the hydrophilic core (Fig. 3, A and B) and should thus be able to accommodate a bulky, N-linked glycan. The side chain of Asn197, however, appears exposed to a quite crowded area of the hydrophilic core (Fig. 3, A and B) and probably do not have enough space to accommodate a bulky, N-linked glycan. Thus, our E196K fibril model would represent the fibril core of mono-glycosylated PrP^Sc^. Very recently, mono-glycosylated PrP^Sc^ has been found to have structural instability (*47*) and form fibrillar plaques (*48*) in some PrP mutations including familial CJD-related mutations V180I and T183A (*48*).

In summary, familial prion disease-related mutation E196K displays a novel amyloid fibril structure revealed by cryo-EM and has a significantly lower conformational stability and protease resistance activity compared to the wild-type fibril. The reported cryo-EM structure of the E196K PrP fibril reveals an unusual overall structure when compared to the wild-type fibril, characterized by a disruption of key salt bridges, a hydrophilic cavity, two unidentified densities flanking the protofibrils and 5 instead of 6 β-strands in the core. The structure provides structural evidence for different prion strains and may inspire future research on the mechanism how the familial mutants of PrP can drive disease, given the issue of polymorphism among amyloid fibrils.

## MATERIALS AND METHODS

### Protein expression and purification

A plasmid-encoding, full-length human PrP (23−231) was a kind gift from Dr. G.-F. Xiao (Wuhan Institute of Virology, Chinese Academy of Sciences). The gene for PrP 23-231 was constructed in the vector pET-30a (+), and a PrP mutant E196K was constructed by site-directed mutagenesis using a wild-type PrP template; the primers are shown in Table S1. All PrP plasmids were transformed into *Escherichia coli*. Recombinant full-length wild-type human PrP and its variant E196K were expressed from the vector pET-30a (+) in *E. coli* BL21 (DE3) cells (Novagen, Merck, Darmstadt, Germany). PrP proteins were purified by high-performance liquid chromatography on a C4 reverse-phase column (Shimadzu, Kyoto, Japan) as described by Bocharova et al (*49*) and Zhou et al (*50*). After purification, recombinant wild-type PrP^C^ and E196K PrP^C^ were dialyzed against 20 mM Tris-HCl buffer (pH 7.4) three times, concentrated, filtered and stored at −80 °C. SDS-PAGE and mass spectrometry were used to confirm that the purified human PrP proteins were single species with an intact disulfide bond. We used a NanoDrop OneC Microvolume UV-Vis Spectrophotometer (Thermo Scientific) to determine the concentrations of wild-type human PrP^C^ and E196K PrP^C^, using their absorbances at 280 nm and the molar extinction coefficients calculated from the composition of the proteins (http://web.expasy.org/protparam/).

### PrP fibril formation

Full-length recombinant wild-type human PrP^C^ and E196K PrP^C^ (40 μM) were incubated in 20 mM Tris-HCl buffer (pH 7.4) containing 2 M guanidine hydrochloride (GdnHCl) with shaking at 180 rpm at 37 °C for 9−11 h and 7−9 h, respectively, and the wild-type and E196K fibrils were collected. Large aggregates in E196K fibril samples and the wild-type fibril samples were removed by centrifugation for 5,000 *g* at 4 °C for 10 min. The supernatants were then dialyzed against 20 mM NaAc buffer (pH 5.0) three times, to ensure that GdnHCl had been removed. After dialysis, E196K fibril samples and the wild-type fibril samples were purified by ultracentrifugation for 100,000 *g* for 30 min twice and washed with 20 mM NaAc buffer (pH 5.0). The pellets containing E196K fibrils or the wild-type fibrils were resuspended in 100 μl of 20 mM NaAc buffer (pH 5.0) and not treated with proteinase K. We used a NanoDrop OneC Microvolume UV-Vis Spectrophotometer (Thermo Scientific) to determine the concentrations of E196K fibril and wild-type PrP fibril, using their absorbances at 280 nm and the molar extinction coefficients calculated from the composition of PrPs (http://web.expasy.org/protparam/).

### Congo red binding assays

E196K fibrils were analyzed by Congo red binding assays. A stock solution of 200 μM Congo red was prepared in PBS and filtered through a filter of 0.22 μm pore size before use. In a typical assay, the E196K fibril sample was mixed with a solution of Congo red to yield a final Congo red concentration of 50 μM and a final PrP concentration of 10 μM, and the absorbance spectrum between 400 and 700 nm was then recorded on a Cytation 3 Cell Imaging Multi-Mode Reader (BioTek).

### TEM of E196K fibrils

E196K fibrils were examined by transmission electron microscopy of negatively stained samples. Ten microliters of E196K fibril samples (∼13 μM) were loaded on copper grids for 30 s and washed with H_2_O for 10 s. Samples on grids were then stained with 2% (w/v) uranyl acetate for 30 s and dried in air at 25 °C. The stained samples were examined using a JEM-1400 Plus transmission electron microscope (JEOL) operating at 100 kV.

### Proteinase K digestion assay

E196K fibril were assessed by proteinase K digestion. E196K fibril samples were incubated with proteinase K at a protease:PrP molar ratio of 1:500 to 1:100 for 1 h at 37 °C. Digestion was stopped by the addition of 2 mM phenylmethylsulfonyl fluoride, and samples were analyzed in 15% SDS-PAGE and detected by silver staining.

### Cryo-EM of PrP fibrils

E196K fibrils were produced as described above. An aliquot of 3.5 μl of ∼13 μM E196K fibril solution was applied to glow-discharged holey carbon grids (Quantifoil Cu R1.2/1.3, 300 mesh), blotted for 3.5 s and plunge-frozen in liquid ethane using an FEI Vitrobot Mark IV. Grids were examined using an FEI Talos F200C microscope, operated at 200 kV and equipped with a field emission gun and a FEI Ceta camera (Thermo Fisher). The cryo-EM micrographs were acquired on a FEI Titan Krios microscope operated at 300 kV (Thermo Fisher) and equipped with a Gatan K2 Summit camera. A total of 4,883 movies were collected in counting mode at a nominal magnification of ×29,000 (pixel size, 1.014 Å) and a dose of 8 e^-^ Å^-2^ s^-1^ (see Table 1). An exposure time of 8 s was used, and the resulting videos were dose-fractionated into 40 frames. A defocus range of −1.5 to −3.0 μm was used.

### Helical reconstruction

All 40 video frames were aligned, summed and dose-weighted by MotionCor2 and further binned to a pixel size of 1.014 Å (*51*). Contrast transfer function estimation of aligned, dose-weighted micrographs was performed by CTFFIND4.1.8 (*52*). Subsequent image-processing steps, include manual picking, particle extraction, 2D classification, 3D classification, 3D refinement and post-processing, were performed by RELION3.1 (*39*).

In total, 25,301 fibrils were picked manually from 4,883 micrographs, and 686 and 400 pixel boxes were used to extract particles by 90% overlap scheme. Two-dimensional classification of 686-box-size particles was used to calculate the initial twist angle. In regard to helical rise, 4.8 Å was used as the initial value. Particles were extracted into 400-box sizes for farther processing. After several iterations of 2D and 3D classification, particles with the same morphology were picked out. Local searches of symmetry in 3D classification were used to determine the final twist angle and rise value. The 3D initial model was built by selected 2D classes; 3D classification was performed several times to generate a proper reference map for 3D refinement. Three-dimensional refinement of the selected 3D classes with appropriate reference was performed to obtain final reconstruction. The final map of E196K fibrils was convergent with a rise of 2.41 Å and twist angle of 179.66°. Post-processing was preformed to sharpen the map with a *B* factor of −43.26 Å^2^. Based on the gold-standard Fourier shell correlation (FSC) = 0.143 criteria, the overall resolution was reported as 3.07 Å. The statistics of cryo-EM data collection and refinement is shown in Table 1.

### Atomic model building and refinement

COOT (*53*) was used to build and modify the atomic model of E196K fibril based on the cryo-EM structure of wild-type PrP fibril (PDB 6LNI) (*33*). The model with 3 adjacent layers was generated for structure refinement. The model was refined using the real-space refinement program in PHENIX (*54*).

### Global denaturation of E196K and wild-type fibrils analyzed by ThT fluorescence and SDS-PAGE

Amyloid fibrils were produced from wild-type and E196K PrPs incubated in 20 mM Tris-HCl buffer (pH 7.4) containing 2 M guanidine hydrochloride and shaking at 37 °C for 10 h. The PrP fibrils were incubated for 1 h at 25 °C in the presence of different concentrations of guanidine thiocyanate (GdnSCN). The concentration of GdnSCN was then adjusted to 0.35 M, followed by a thioflavin T (ThT) binding assay. A Cytation 3 Cell Imaging Multi-Mode Reader (BioTek) was used to the ThT fluorescence produced, with excitation at 450 nm and emission at 480 nm. The half-concentration at which the ThT fluorescence intensity of PrP fibrils is decreased by 50% (*C*_1⁄2_) of E196K fibril and the wild-type fibril were determined using a sigmoidal equation (*55, 56*) using the above ThT fluorescence data. The PrP fibrils were also dialyzed against 20 mM NaAc buffer (pH 5.0) for three times and diluted to a final concentration of 16 μM using 10% SDS. The final concentration SDS was 6%. The PrP fibrils were incubated with 6% SDS for 5 min under a thermal gradient from 25−100 °C, mixed with the 5 × loading buffer (without SDS, β-mercaptoethanol and heating) and separated by 12.5% SDS-PAGE. The soluble PrP monomers were detected by SDS-PAGE with Coomassie Blue R250 staining.

## Acknowledgments

Cryo-EM data was collected at the Center of Cryo Electron Microscopy, Zhejiang University, China. We thank G.-F. Xiao (Wuhan Institute of Virology, Chinese Academy of Sciences) for the kind gift of the human PrP^C^ plasmid; S. Chang (Center of Cryo Electron Microscopy, Zhejiang University), L. Wu (Center of Cryo Electron Microscopy, Zhejiang University) and X. Zhang (Zhejiang University School of Medicine) for their technical assistance with Cryo-EM; and Y. Wang (Institute of Biophysics, Chinese Academy of Sciences) for her helpful suggestions.

## Author contributions

P.Y., C.L. and Y.L. supervised the project. L.-Q.W., C.L. and Y.L. designed the experiments. L.-Q.W., H.-Y.Y., X.-N.L. and J.C. purified the E196K PrP^C^ and E196K fibrils. L.-Q.W., H.-Y.Y. and H.-B.D. performed Congo red binding, proteinase K digestion and conformational stability assays of PrP fibrils. L.-Q.W., K.Z., H.-Y.Y., Y.M., Q.W., C.W., Y.S. D.Z. and D.L. collected, processed and/or analyzed cryo-EM data. L.-Q.W., K.Z., C.L. and Y.L. wrote the manuscript. All authors proofread and approved the manuscript.

## Data and materials availability

All data needed to evaluate the conclusions in the paper are present in the paper and/or the Supplementary Materials. Cryo-EM density maps and the atomic model of human E196K PrP fibrils are available through the Electron Microscopy Data Bank and Protein Data Bank with accession codes EMD-30887 and PDB 7DWV, respectively. Additional data related to this paper may be requested from the authors.

## Supplementary Materials

**Table S1.**
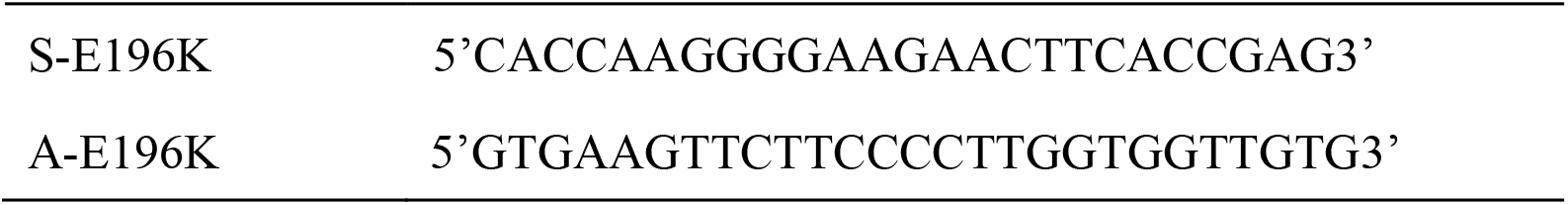
The primers designed for full-length human PrP with E196K mutation.

**Figure S1.**
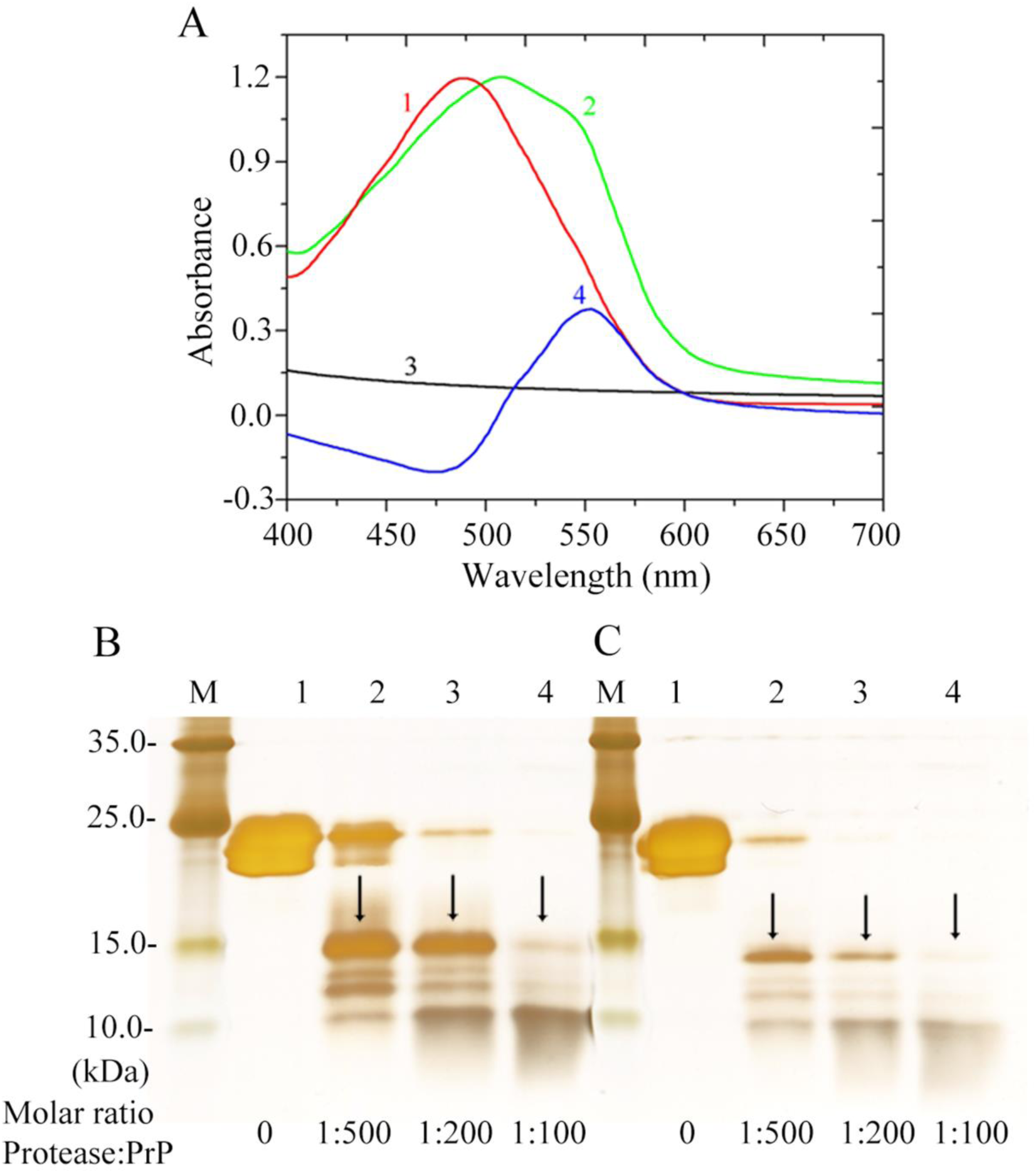
E196K fibrils bind to Congo red and are proteinase K resistant. (**A**) Amyloid fibrils of full-length human PrP with E196K mutation analyzed by Congo red binding assays. The difference spectra (Curve 4, blue) with the maximum absorbance at 550 nm were obtained by subtracting the absorbance spectra of E196K fibrils alone (Curve 3, black) and Congo red alone (Curve 1, red) with the maximum absorbance at 490 nm from those of E196K fibrils + Congo red (Curve 2, green). Congo red binding assays were carried out at 37 °C. (**B** and **C)**, Amyloid fibrils of full-length wild-type human PrP (**B**) and its variant E196K (**C**) analyzed by concentration-dependent proteinase K digestion assays (*33, 50*). Protease-resistant core fragments of 15−16-kDa (**B**) and 14−15-kDa (**C**) are highlighted using black arrows. Samples were treated with proteinase K for 1 h at 37 °C at protease:PrP molar ratios of 1:500 (lane 2), 1:200 (lane 3) and 1:100 (lane 4). The control with no protease was loaded in lane 1. Molecular weight markers were loaded on lane M. Protein fragments were separated by SDS-PAGE and detected by silver staining. These experiments were repeated three times with different batches of fibrils and similar results.

**Figure S2.**
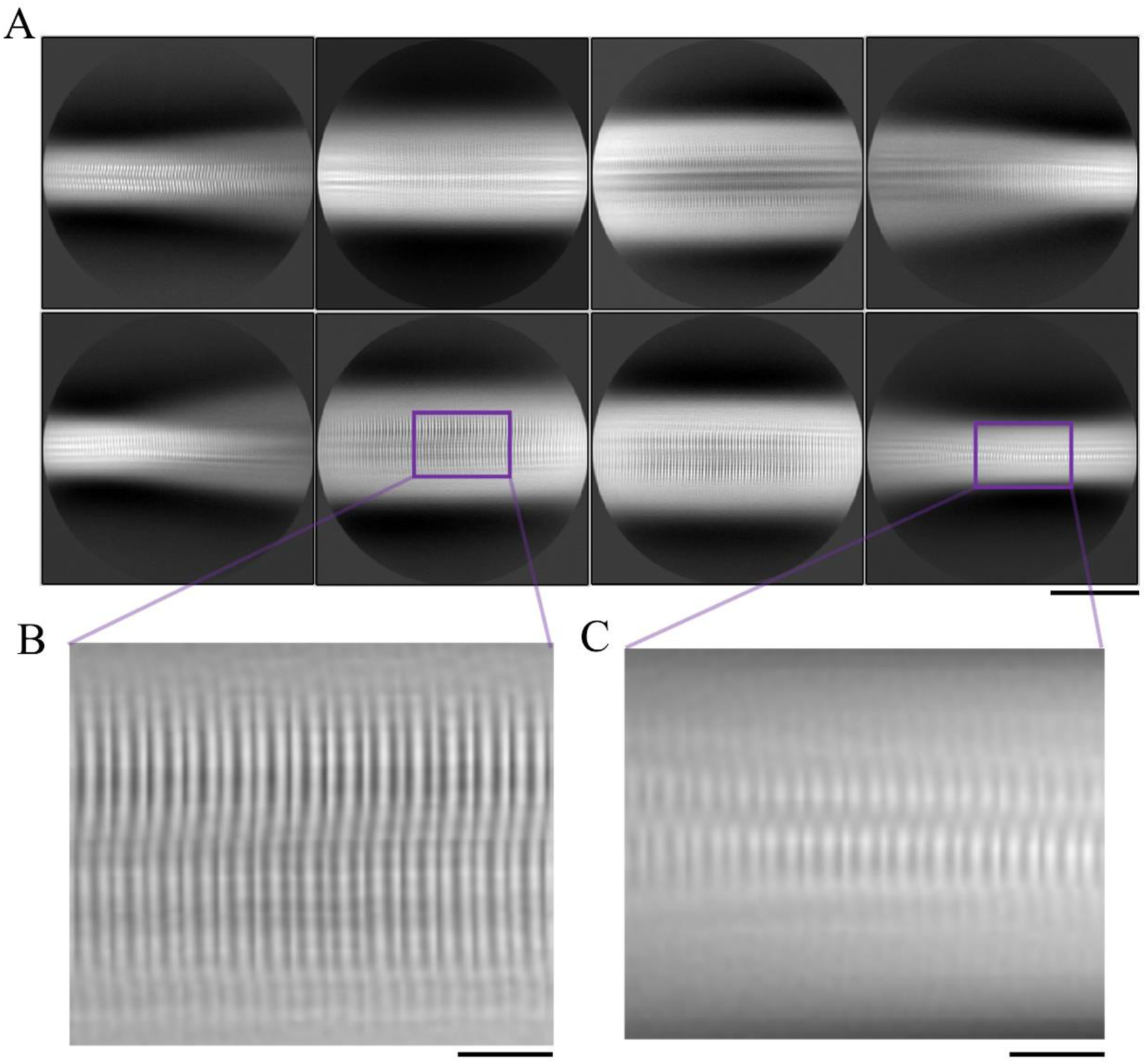
Cryo-EM images of E196K fibrils. (**A**) Reference-free 2D class averages of E196K fibrils showing two protofibrils intertwined. Scale bar, 10 nm. (**B** and **C)** Enlarged images of (**A**) showing two protofibrils arranged in a staggered manner. Scale bar, 2 nm.

**Figure S3.**
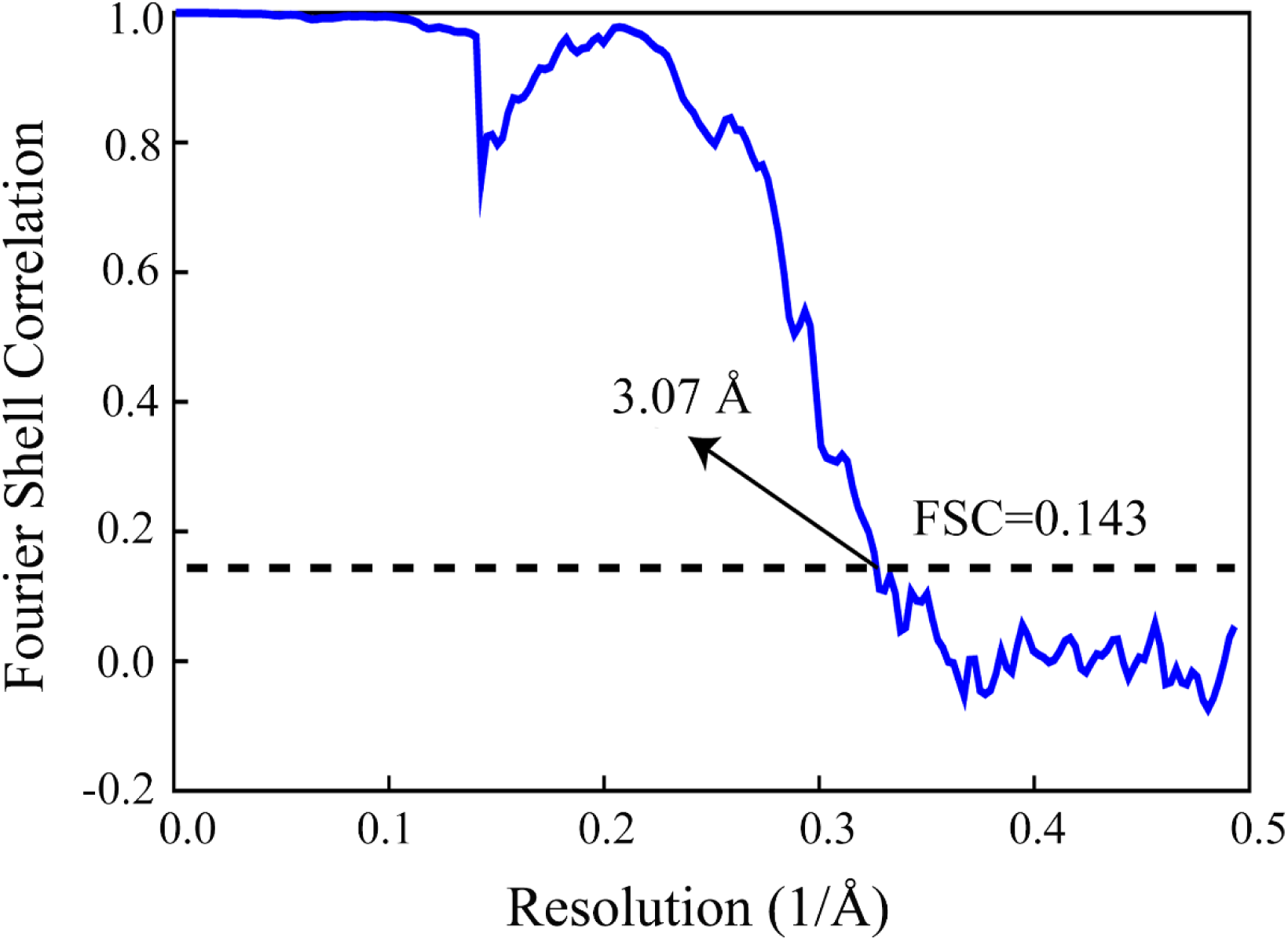
Global resolution estimate for the E196K fibril reconstructions. Gold-standard refinement was used for estimation of the density map resolution. The global resolution of 3.07 Å was calculated using a Fourier shell correlation (FSC) curve cut-off at 0.143.

**Figure S4.**
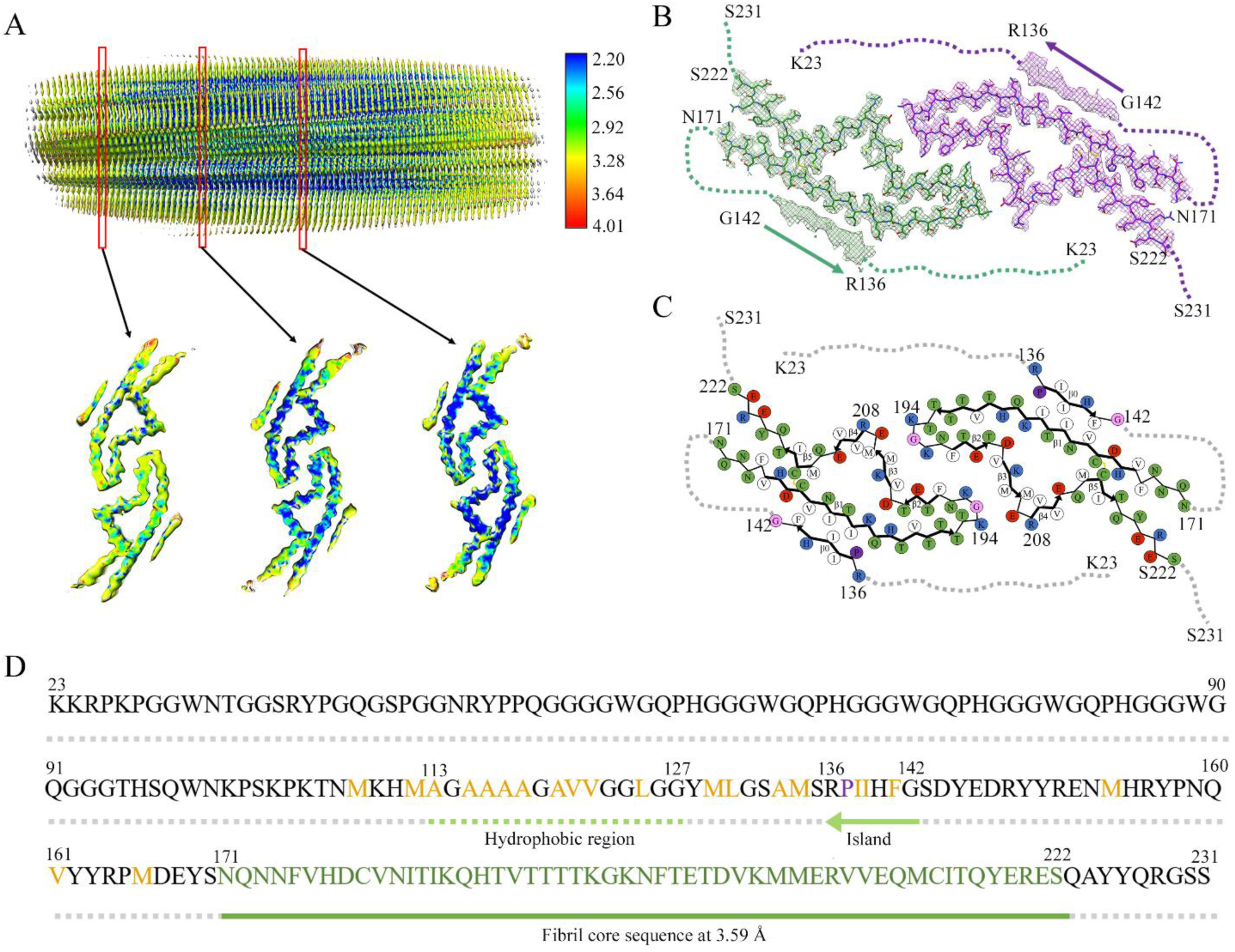
Cryo-EM structure of E196K fibrils at 3.59 Å. (**A**) Local resolution estimate for the E196K fibril reconstructions at 3.59 Å. The unmasked density map of E196K fibrils is colored according to local resolution estimated by ResMap. The three enlarged cross sections show the left top view of the density map of two protofibrils, and each monomer has an island. The color key on the right shows the local structural resolution in angstroms (Å) and the colored map indicates the local resolution ranging from 2.2 to 4.01 Å. (**B**) Cryo-EM map of E196K fibrils at 3.59 Å with a fibril core comprising residues 171−222 from E196K. Two protofibrils with two islands (marked by arrow) are colored green and purple, respectively. Each island, probably comprising residues 136−142 from E196K and forming a β-strand (β0), is located on the opposing side of hydrophobic side chains of Val180, Ile182 and Ile184 in each monomer. The side chains of Val180, Ile182, Ile184, Pro137, Ile139 and Phe141 forms a hydrophobic steric zipper-like interface, thereby stabilizing the E196K fibrils. (**C**) Schematic view of E196K fibril core at 3.59 Å. Residues are colored as follows: white, hydrophobic; green, polar; red and blue, negatively and positively charged, respectively; purple, proline; magenta, glycine. Six β-strands (β0-b5) are indicated with bold lines. E196K fibrils are stabilized by a disulfide bond (yellow line) formed between Cys179 and Cys214 in each monomer, also visible in panel **B**. (**D**) The primary sequence of E196K, in which the green bar marks E196K fibril core comprising residues 171−222 at 3.59 Å. The green arrow marks an island flanking each protofibril, and the green dashed line marks a hydrophobic region of human PrP^C^. Residues are colored as follows: orange, hydrophobic; green, E196K fibril core at 3.59 Å. The dashed lines correspond to residues (23−135)/ (143−170)/ (223−231) not modeled in the cryo-EM density.

**Figure S5.**
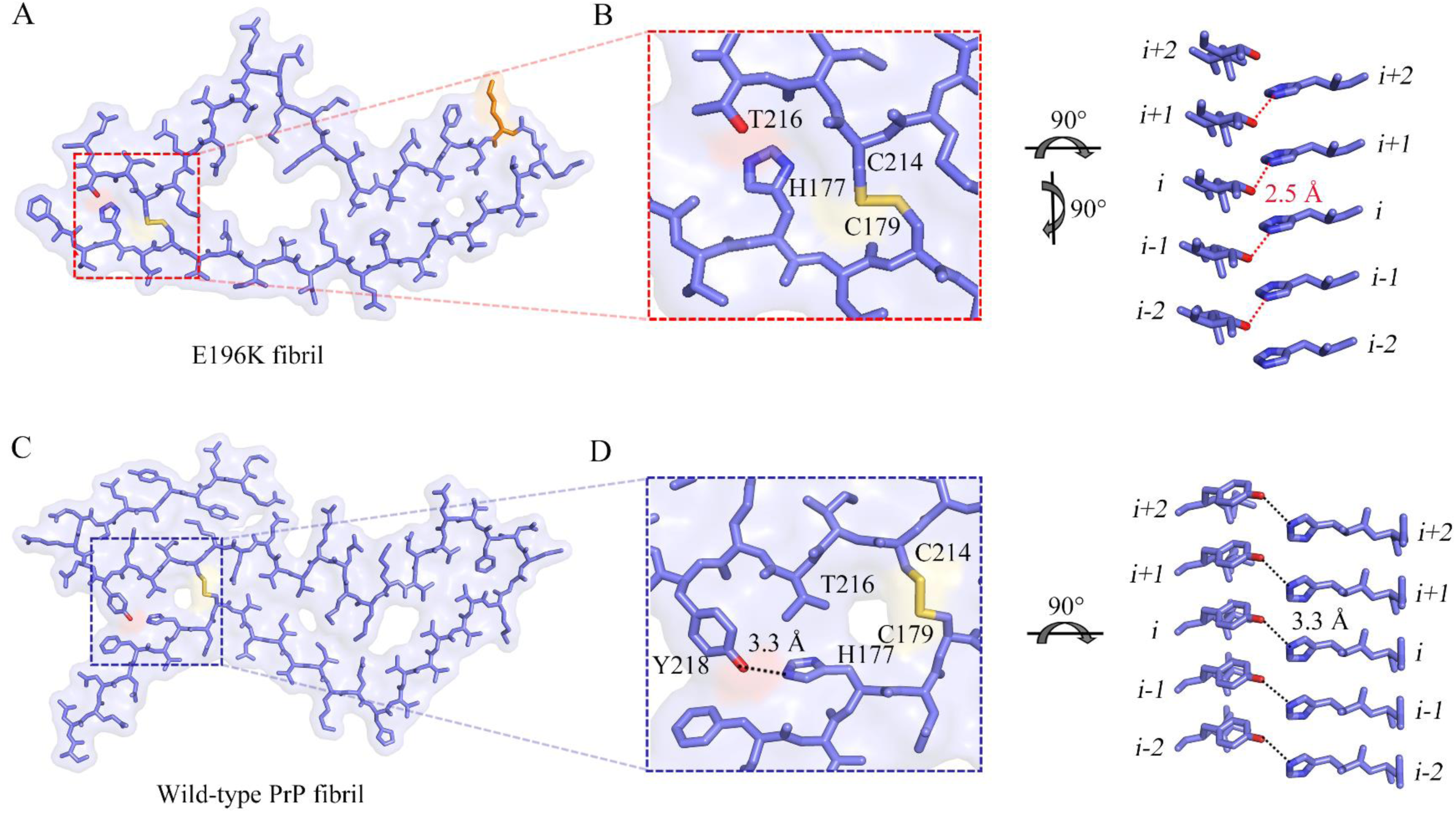
Close-up view of the stick representation of the structures of E196K and wild-type fibrils. (**A**) A space-filled model overlaid onto a stick representation of E196K fibril in which one protofibril is shown in blue. An E196K protofibril is stabilized by one disulfide bond (yellow line) between Cys179 and Cys214, and the disulfide bond region is magnified in **B**. Lys196 in E196K variant is highlighted in orange. **(B**) A magnified top view of the disulfide bond region of an E196K protofibril, where a disulfide bond (yellow line) is formed between Cys179 and Cys214 and a hydrogen bond is formed between His177 and Thr216. A side view (right) highlighting a hydrogen bond between His177 (*i*) and Thr216 from its opposing adjacent subunit (*i-1*), with a distance of 2.5 Å (red). (**C**) A space-filled model overlaid onto a stick representation of wild-type PrP fibril (PDB 6LNI) (*33*) in which one protofibril is shown in blue. A wild-type PrP protofibril is also stabilized by one disulfide bond (yellow line) between Cys179 and Cys214, and the disulfide bond region is magnified in **D**. (**D**) A magnified top view of the disulfide bond region of a wild-type PrP protofibril, where a disulfide bond (yellow line) is formed between Cys179 and Cys214 and a hydrogen bond is formed between His177 and Tyr218. A side view (right) highlighting a hydrogen bond between His177 (*i*) and Tyr218 from its opposing subunit (*i*), with a distance of 3.3 Å (black).

**Figure S6.**
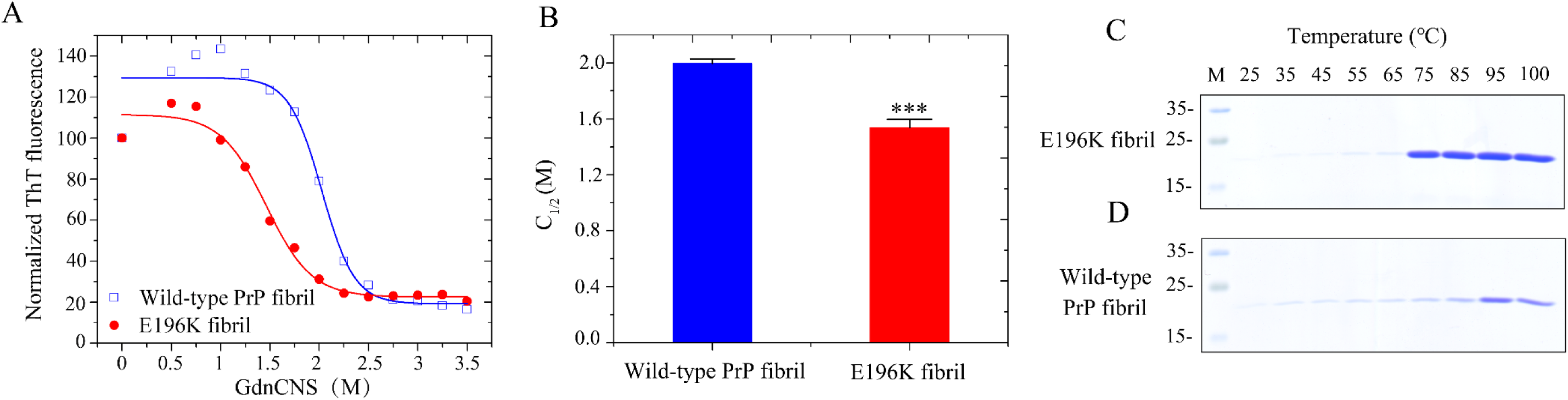
The E196K mutation significantly decreases the conformational stability of PrP fibrils. Amyloid fibrils were produced from wild-type and E196K PrPs incubated in 20 mM Tris-HCl buffer (pH 7.4) containing 2 M guanidine hydrochloride and shaking at 37 °C for 10 h. (**A**) GdnSCN-induced denaturation profiles were monitored for E196K fibril (red) and wild-type PrP fibril (blue). The PrP fibrils were incubated for 1 h at 25 °C in the presence of different concentrations of GdnSCN. The concentration of GdnSCN was then adjusted to 0.35 M, followed by a ThT binding assay. The solid lines show the best sigmoidal fit for the ThT intensity-time curves. (**B**) The *C*_1⁄2_ (the half-concentration at which the ThT fluorescence intensity of PrP fibrils is decreased by 50%) values of E196K fibril (red) and wild-type PrP fibril (blue) were determined using a sigmoidal equation (*55, 56*) using the ThT fluorescence data obtained, and are expressed as mean ± SD of the values obtained in 4 independent experiments. E196K fibril, *p* = 0.0000094. The revised Student *t* test was used to perform statistical analyses. Values of *p* < 0.005 indicate statistically significant differences. The following notation is used throughout: *, *p* < 0.005; **, *p* < 0.001; and ***, *p* < 0.0001 relative to wild-type PrP fibril (a control). (**C** and **D**) Thermal stabilities of E196K fibril (**C**) and wild-type PrP fibril (**D**) under different temperatures were determined by SDS-PAGE. The PrP fibrils were incubated with 6% SDS for 5 min under each temperature indicated, and the soluble PrP monomers were detected by SDS-PAGE with Coomassie Blue R250 staining. The melting temperature (*T*_m_) values of E196K fibril (**C**) and wild-type PrP fibril (**D**) were ∼75 °C and ∼95 °C, respectively, at which the band intensity of soluble PrP monomers reached a maximum. Molecular weight markers were loaded on lane M. The E196K fibril has a significantly lower conformational stability compared to the wild type. These experiments were repeated three times with different batches of fibrils and similar results.

